# Information transfer from spatial to social distance in rats: implications for the role of the posterior parietal cortex in spatial-social integration

**DOI:** 10.1101/2024.10.14.618305

**Authors:** Taylor B. Wise, Victoria Templer, Rebecca D. Burwell

## Abstract

Humans and other social animals can represent and navigate complex networks of social relationships in ways that are suggestive of representation and navigation in space. There is some evidence that cortical regions initially required for processing space have been adapted to include processing of social information. One candidate region for supporting both spatial and social information processing is the posterior parietal cortex (PPC). We examined the hypothesis that rats can transfer or generalize distance information across spatial and social domains and that this phenomenon requires the PPC. In a novel apparatus, rats learned to discriminate two conspecifics positioned at different spatial distances (near vs. far) in a goal-driven paradigm.

Following spatial learning, subjects were tested on probe trials in which spatial distance was replaced with social distance (cagemate vs. less familiar conspecific). The PPC was chemogenetically inactivated during a subset of probe sessions. We predicted that, in control probe trials, subjects would select conspecifics whose social distance matched the previously learned spatial distance. That is, if trained on the near distance, the rat would choose the highly familiar cagemate, and if trained on the far distance, the rat would choose the less familiar conspecific. Subjects learned to discriminate conspecifics based on spatial distance in our goal-driven paradigm. Moreover, choice for the appropriate social distance in the first probe session was significantly higher than chance. This result suggests that rats transferred learned spatial information to social contexts. Contrary to our predictions, PPC inactivation did not impair spatial to social information transfer. Possible reasons are discussed. To our knowledge, this is the first study to provide evidence that spatial and social distance are processed by shared cognitive mechanisms in the rat model.

## INTRODUCTION

Processing multiple forms of information is required for engaging with complex environments. Two information types that appear distinct, but are necessary for daily life, are spatial and social information. Cognitive and neuroscience research suggests that these domains intersect in our larger understanding of the world. Yet, there are open questions about the mechanisms that contribute to spatial and social cognition and how they may be shared across domains.

One way spatial and social cognition may intersect is through broad cognitive mapping skills that exist on a spectrum of physical and abstract space (Figure 1). At one end, cognitive maps can be used to navigate physical space (Tolman, 1948) and are confirmed in the brain through specialized neurons such as place cells (O’keefe & Nadel, 1978), grid cells (Hafting, Fyhn, Molden, Moser, & Moser, 2005; Stensola et al., 2012), and many others (Grieves & Jeffery, 2017). These cells are context dependent and can be remapped for flexible decision making (Alexander et al., 2016; Colgin, Moser, & Moser, 2008). They also reflect rewarding locations during goal-oriented spatial navigation (Ormond & O’Keefe, 2022). Abstract information, such as social information, may also be encountered in physical spaces, such as when two individuals cross paths on a walkway. Recent studies on social place cells – neurons that selectively fire when a social partner occupies a specific location – support this form of cognitive mapping in non-human animals (Danjo, Toyoizumi, & Fujisawa, 2018; Omer, Maimon, Las, & Ulanovsky, 2018). Through cognitive maps of physical space, spatial distance from environmental landmarks or from social entities can be estimated. At a higher level of abstraction, cognitive maps may also aid in processing social information independent of physical space. These abstract maps are defined by the strength of social relationships, or social distance, between oneself and others. While studies investigating social cognitive maps are relatively new, there is striking evidence for systematic mapping of social space in the hippocampal formation (Montagrin, Saiote, & Schiller, 2018; Schafer & Schiller, 2018; Tavares et al., 2015), a collection of brain regions widely associated with encoding cognitive maps (Geva-Sagiv, Las, Yovel, & Ulanovsky, 2015; Hunsaker & Kesner, 2018). Neural correlates for other abstract concepts, such as reward and time, further support the existence of non-physical cognitive maps (Eichenbaum, 2014; Knudsen & Wallis, 2021; Schiller et al., 2015).

**Figure 1.**
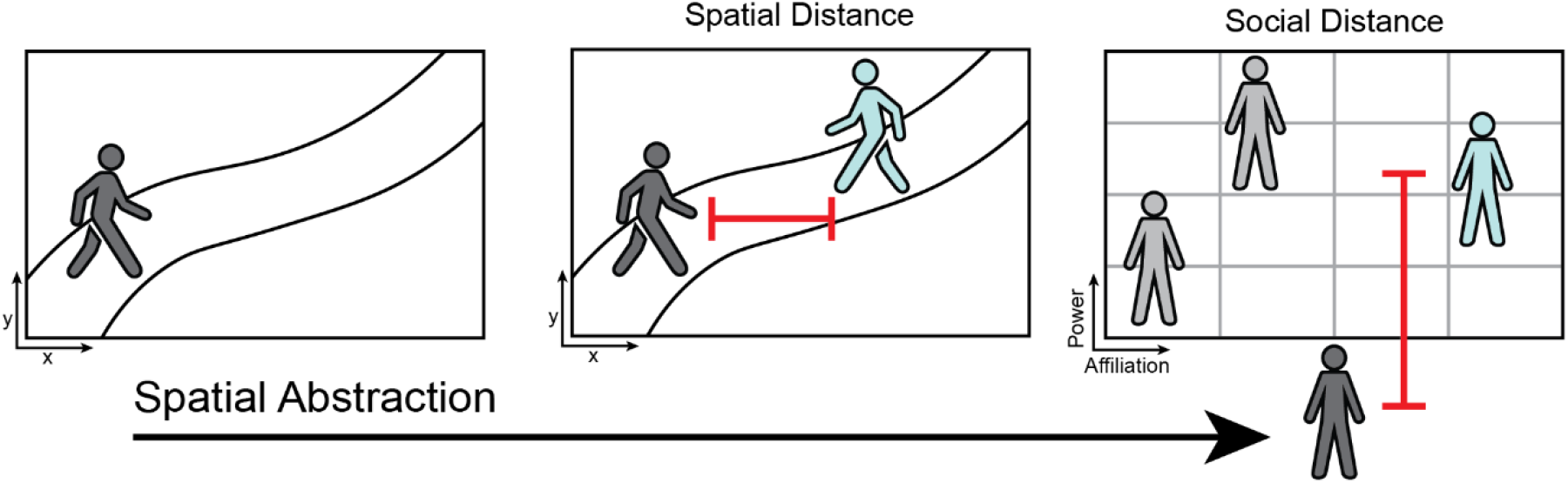
Spatial abstraction. Physical space (left) includes one’s position in an environment that can be used to build a spatial cognitive map. Physical space with social information (middle) includes the position of oneself and other social actors within an environment. Spatial distance between oneself and others (red line) can be estimated and contribute to spatial cognitive maps. Social space (right) includes abstract representations of social dynamics (e.g. power and affiliation) between oneself and others. Social distance between oneself and others (red line) can be estimated and contribute to social decision making and abstract cognitive maps. As environments increase in abstraction individuals may continue to process information similarly. Adapted from Schafer & Schiller (2018).

Overall, prior research suggests that maintaining cognitive maps is crucial for adaptively navigating spatial and social environments. Moreover, cross-species evidence for cognitive mapping may point to shared evolutionary traits in social mammals.

Evolutionary theories state that as social interaction became increasingly important for survival, the neocortex was altered to meet these complex cognitive demands via exaptation (Parkinson & Wheatley, 2013). Exaptation is the evolutionary process in which preexisting traits are co-opted to meet new and emerging demands (Gould & Vrba, 1982). In the case of spatial-to-social exaptation, cortical regions initially required for spatial processing may have been later recruited for social processing. One area proposed to have been co-opted for social cognition is the posterior parietal cortex (PPC) (Parkinson & Wheatley, 2013). A rich literature substantiates the role of PPC in spatial cognition across human and non-human primates (Freedman & Ibos, 2018; Sack, 2009) and rodents (Kesner, 2009; Kesner & Creem-Regehr, 2013). The PPC is also thought to aid in processing abstract cognition (Hubbard, Piazza, Pinel, & Dehaene, 2005; Yamazaki, Hashimoto, & Iriki, 2009), including social cognition (Fujii, Hihara, & Iriki, 2008; Parkinson & Wheatley, 2013), though this literature is less extensive. Interestingly, abstract processing in the PPC is observed when task demands require integration of abstract and spatial information. The majority of these studies focus on PPC contributions to spatial-temporal integration (Hubbard et al., 2005; Oliveri et al., 2009; Riemer, Diersch, Bublatzky, & Wolbers, 2016). Despite compelling evidence for the role of the PPC in integrating spatial and abstract information, few studies have directly addressed its contributions to integrating spatial and social information. Limited evidence shows that the human PPC is recruited during both spatial and social distance judgements (Giardina, Caltagirone, Cipolotti, & Oliveri, 2012; Parkinson, Liu, & Wheatley, 2014; Yamakawa, Kanai, Matsumura, & Naito, 2009). In (Yamakawa et al., 2009) fMRI participants separately estimated spatial and social ego-centric distance and the PPC was similarly activated in both conditions. In a second experiment, participants spontaneously placed figurines of themselves and social partners previously experienced in the scanner on a physical map. Researchers found that physical distance between self- and partner-figurines matched the social distance previously reported during fMRI scans, suggesting spatial-social integration. Brain activity was not recorded during the physical distance component of the figurine task. However, other research suggests that the right inferior parietal lobule (IPL), a section of the PPC, may encode spatial and social distance (Parkinson et al., 2014). While these studies provide interesting evidence for shared spatial and social mechanisms in the PPC, similar questions have yet to be investigated in non-human animals.

Understanding the neural interplay between spatial and social cognition using animal models will necessarily require experimental designs that measure observable behavior rather than verbal self-reports. One possibility for evaluating shared spatial-social mechanisms is to test whether an animal’s demonstrated knowledge of a spatial distance can be applied to a social distance context. Rats are a suitable option for this approach given their spatial prowess and social nature. They are highly skilled navigators attuned to many features of the environment, including distance. Rats can detect changes in the spatial distance and position of environmental landmarks and can discriminate towards spatial novelty (Goodrich-Hunsaker, Hunsaker, & Kesner, 2005; Save, Poucet, Foreman, & Buhot, 1992; Warburton & Brown, 2015). Spatial discrimination tasks often examine how rats move through the environment to interact with landmarks or other spatial cues. Indeed, rats benefit from motion cues in these tasks. Demands for discriminating spatial distance without movement are constrained by rats’ forward-facing depth perception, low acuity, and dependence on motion cues (Burn, 2008; Legg & Lambert, 1990; Powers & Green, 1978) . Rats can, however, discriminate some static visual patterns including patterns and shapes. Moreover, visual acuity in rats is higher in the lower half of their visual field (Lashley, 1932, 1938; Minini & Jeffery, 2006), and this can be leveraged to discriminate stimuli presented on the floor (Furtak et al., 2009). One way to strengthen such discrimination tasks would be to maximize the use of multimodal information. Rats are also skilled at detecting differences in social distances, primarily by discriminating towards social novelty (Acikgoz, Dalkiran, & Dayi, 2022; Templer, Wise, Dayaw, & Dayaw, 2018). Even when novelty is equalized, rats distinguish individual conspecifics (Gheusi, Goodall, & Dantzer, 1997). A number of spatial discrimination paradigms, and the majority of social discrimination paradigms, measure changes in spontaneous and unstructured exploration of stimuli. While these studies are critical for evaluating naturalistic rat behavior, it would be useful to have reward-based spatial or social distance tests. A reward-based discrimination task would allow for more extended training and provide more reliable assessment of behavior. While there are some reward-based tasks for spatial cognition, to our knowledge, none require discrimination of spatial distance (though see (Klement, Levcik, Duskova, & Nekovarova, 2010). Discrimination of social distance on the other hand is regularly studied (Acikgoz et al., 2022) but has only once been paired with a reward-based task (Gheusi et al., 1997). Crucially, all of these tests have limitations for examining interactions between spatial and social information processing. What is needed is a goal-oriented paradigm that is suitable for both domains. Rats are well suited for complex decision-making tasks in both spatial (Scott et al., 2021; van der Staay, Gieling, Pinzón, Nordquist, & Ohl, 2012) and non-spatial domains (Izquierdo & Belcher, 2012; Robbins, 2002; Tait, Bowman, Neuwirth, & Brown, 2018), though very little has been dedicated to social processing and spatial-social integration. A necessary step towards exploring the neural bases of putative shared spatial-social mechanisms in rats will be to design tasks with similar demands for processing spatial and social information, particularly for goal-oriented distance discrimination.

In this study, we examine 1) whether learned spatial information can be transferred to a socially relevant context using a spatial distance/social distance paradigm and 2) whether the PPC is required for such transfer. Rats were trained to discriminate spatial distance from a conspecific in a goal-oriented task and then tested on social distance probes. If rats are capable of transferring information across domains, then during probes subjects will select social distances that match previously learned spatial distances. We used a novel, vertical maze task that required reliance on auditory and olfactory cues as well as visual cues. To test whether the PPC is required for potential spatial-to-social transfers, designer receptors exclusively activated by designer drugs (DREADDs) were used to temporarily inactivate the dorsal PPC during probe trials. If the PPC is required for spatial to social domain transfer, then rats with PPC inactivation will be impaired in accurately discriminating social distances. To our knowledge, this is the first investigation of 1) shared spatial-social mechanisms within a single paradigm in rats, and 2) PPC function for spatial-social domain transfer. Our results contribute to understanding how spatial and social cognition interact and offer valuable insight into cognitive mapping research. These findings also highlight rats’ ability to process complex information in a goal-directed decision-making paradigm. Finally, these results aid in determining when the PPC may be required in spatial and social processing.

## METHODS

### Subjects

Twenty-four adult male Long Evans rats were used in this study (Charles River Laboratories, Boston, MA, USA). Twelve rats were used as subjects and 12 rats were used as non-subject demonstrators. Subjects and demonstrators had no social interaction outside of testing and demonstrators were equally unfamiliar to subjects. Demonstrators remained the same across test phases. All rats were pair-housed in a ventilated cage rack system and were given ad libitum access to food until reaching 85- 90% of free feeding weight and were then diet restricted. Individuals’ weight was monitored weekly to ensure comparable size between cagemates and assigned demonstrators. Access to water was unrestricted. Rats were placed on a 12:12 reversed light-dark cycle to ensure all testing was conducted at a consistent time during the dark phase. Prior to any testing or surgery, rats were well-handled by experimenters. Surgery was conducted when rats were approximately 3 months of age and weighed 300-400 grams. All procedures were conducted according to the NIH guidelines for the care and use of rats in research and methods were approved by the Institutional Animal Care and Use Committee (IACUC) at Brown University.

### Surgery

Subjects were divided into sham (n=4) and virus (n=8) groups prior to surgery. Preoperatively, rats were administered diazepam (2.5 mg/kg) intraperitoneally and glycopyrrolate (0.05 mg/kg), buprenorphine (0.03 mg/kg), subcutaneously. Scalps were shaved to expose the surgical site and injected with lidocaine (4 mg/kg) before incisions. Once fully anesthetized with isoflurane rats were secured in a stereotaxic frame such that bregma and lambda were in the same horizontal plane (±0.2 mm).

Craniotomies were achieved with a dental drill and dura was removed. For the virus group, bilateral injections of the DREADD virus AAV8-CamKIIa-hM4d(Gi)-mCherry were administered via pressure injection using a 26-gauge, 0.5 mL capacity Hamilton® syringe (Sigma-Aldrich) at 0.5 μl/min for 5 minutes per site. The syringe remained in position at each site for an additional 5 minutes and was then slowly retracted. Three sites were targeted bilaterally (PPC1-3) at a 0° angle. Anterior to posterior (AP) and medial to lateral (ML) coordinates were calculated relative to bregma, and dorsal to ventral (DV) coordinates were calculated relative to the surface of the cortex. The stereotaxic coordinates (AP, ML, DV) were developed using Olsen and Witter (2016) and Paxinos and Watson (2013), and adjusted for virus spread in pilot animals (Table 1). For the sham group, rats received craniotomies and dura mater removal only. At the end of surgery, the scalp was sutured and post-operative, subcutaneous injections of meloxicam (4 mg/kg, sustained release) and saline (6 mL) were administered. Pair-housed subjects underwent surgery within 1-3 days of each other to minimize separation during initial recovery. Pair-housed subjects were reintroduced in a neutral environment once there was no threat of harm to the surgical site, typically 3-5 days post-op. No aggressive behavior between reintroduced cagemates was observed. All subjects were given at minimum two weeks to recover before proceeding with behavioral testing. No surgery was conducted on demonstrator rats.

**Table 1.**
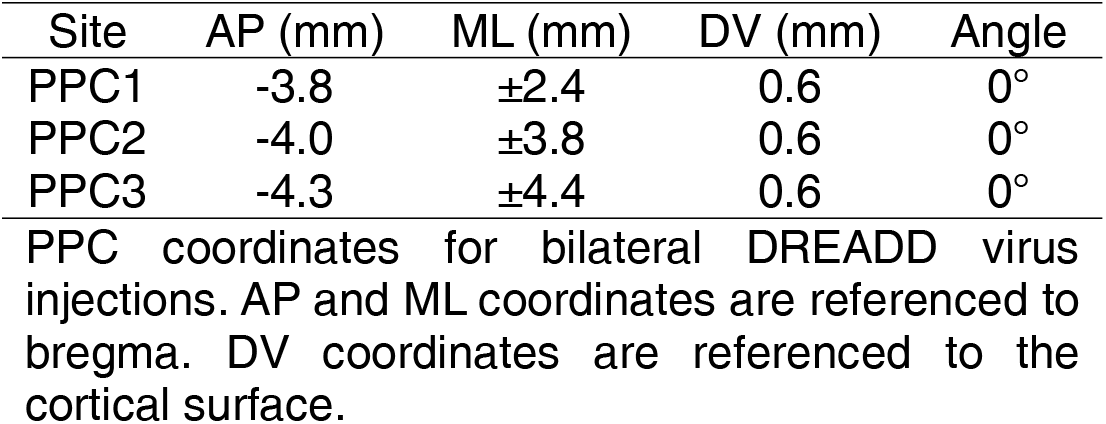
Stereotaxic coordinates for the PPC

| Site | AP (mm) | ML (mm) | DV (mm) | Angle |
| --- | --- | --- | --- | --- |
| PPC1 | -3.8 | ±2.4 | 0.6 | 0° |
| PPC2 | -4.0 | ±3.8 | 0.6 | 0° |
| PPC3 | -4.3 | ±4.4 | 0.6 | 0° |

PPC coordinates for bilateral DREADD virus injections. AP and ML coordinates are referenced to bregma. DV coordinates are referenced to the cortical surface.

### Apparatus

Our novel apparatus, the Vertical Maze (VM), was used to test spatial and social discriminations within one environment (Figure 2). The vertical design of this apparatus allows for ecologically relevant exploration of stimuli presented underfoot. Rich social stimuli promotes processing outside of purely visual information, though visual perception is enhanced by vertical as opposed to front-facing stimuli (Burn, 2008; Furtak et al., 2009; Jeffery & Anderson, 2003; Lashley, 1938; Minini & Jeffery, 2006). The VM included two vertical columns (33 x 52 x 87.6 cm, each) and a three-chamber testing platform with solid Plexiglass walls (91.4 x 52 x 61 cm). Vertical columns were designed to hold plastic rat cages, stated hereby as holding cages, at three distinct spatial distances (0, 58.4, and 87.6 cm) from the testing platform. Holding cages with access holes on the bottom and lid were stacked in each vertical column for subject vertical exploration. Holding cages with solid bottoms and wire bar lids were used to house demonstrator rats during testing and included bedding and chew toys. Vertical columns were constructed of T-slotted aluminum framing and corrugated plastic sheets with sliding privacy doors to create a dark, quiet environment for demonstrators. Testing chambers included modular floors made of aluminum, clear plexiglass with access holes, and wire grid for lever training, vertical exploration, and testing, respectively.

**Figure 2.**
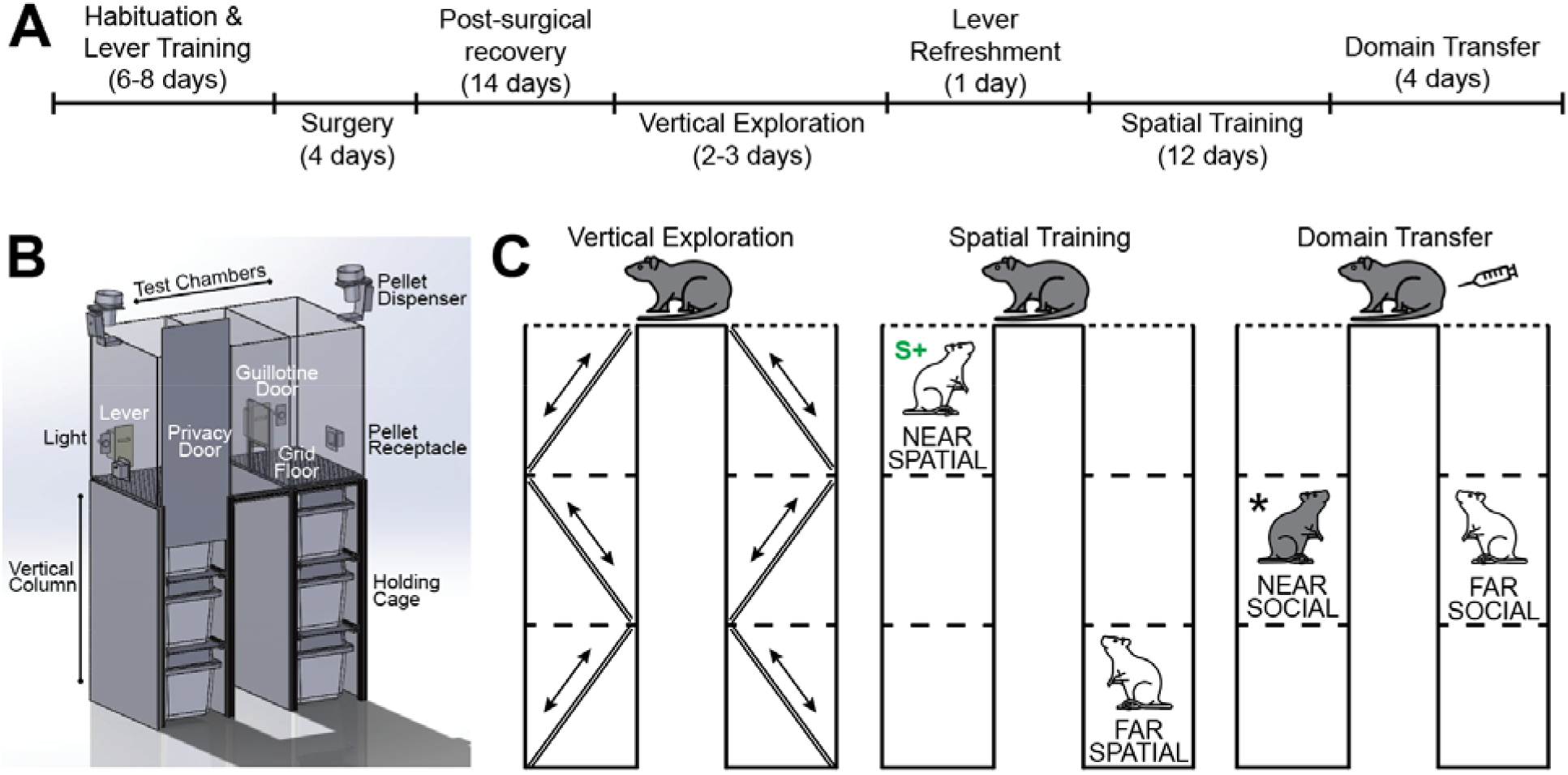
Experimental procedures. A) Timeline of all surgical and behavioral procedures. B) Illustration on the Vertical Maze (VM) apparatus. Automated hardware includes levers, lights, pellet dispensers, pellet receptacles, and guillotine doors. Modular floors were used in the test chamber for each test phase, including grid floors (pictured) for spatial training and domain transfer testing. Holding cages could be placed at the near, middle, or far position in each vertical column relative to test chambers. C) Behavioral procedures in the VM. Vertical exploration allowed subjects (gray) to explore all levels of the vertical columns and test chambers using a ladder system. Spatial training required rats to press a lever for food reward in either the West or East test chamber where they could perceive a demonstrator underneath at a near and far spatial distance. Rewarded choice was counterbalanced across subjects for near- (pictured) and far-spatial distance. Domain transfer testing included trials identical to spatial training as well as social transfer probes. During social transfer probes demonstrators were placed in the middle position to remove differences in spatial distance. One demonstrator was replaced with the subject’s cagemate to create a difference in social distance. Intraperitoneal injections of CNO or saline were conducted prior to each domain transfer testing session. * indicates the correct match between trained spatial distance discrimination and probed social distance discrimination.

Flooring in the middle (start) chamber was always solid plexiglass. Hardware in each testing chamber included an automated lever, light, speaker, pellet dispenser, and pellet receptacle (Med Associates, Inc.), as well as a custom guillotine door to control subject access to adjacent chambers. The VM used a computer interface for controlling all hardware via Ethovision XT (Noldus Information Technology) and tracking all behavior. A high-speed video camera was positioned above the VM and provided real-time feedback to tracking software for behavior-based hardware control. Further details on the VM can be found in Wise, Templer, and Burwell (2024).

### Behavioral Procedures

#### Habituation, Lever Training, & Vertical Exploration

Prior to behavioral testing all rats were habituated over multiple sessions to the VM. Demonstrator rats were habituated to holding cages for 30 minutes a day for two days. Holding cages were placed in either the West or East vertical column and equipped with bedding, enrichment toys, and food pellets. Automated hardware was intermittently turned on so that demonstrators grew accustomed to all maze noises before testing. Subjects were habituated to the testing chambers with an aluminum floor, but without automation, for 10 minutes a day for two days. Guillotine doors remained open to allow for exploration of the West, center, and East chamber. Precision pellets (Bio-Serv©) were scattered throughout chambers and in the West and East pellet receptacles to encourage exploration. No signs of distress, such as freezing or excess grooming, were observed for any rats by the end of habituation.

Subjects were next trained to press a lever for reward that could later be used to indicate choice during testing. Rats were constrained to one test chamber at a time and did not have access to vertical columns. Magazine training consisted of 15-minute sessions in which a pellet was dispensed into the receptacle every 30 seconds. No lever was present during magazine training. Criteria to move on to the next phase of lever training was retrieval of each pellet within 10 seconds for two consecutive days.

Rats were then hand-shaped to press a lever for a pellet. Sessions were limited to one test chamber at a time. The experimenter, watching from the control room, could dispense pellets remotely. Subjects were initially rewarded with a pellet for approaching the lever, and progressively shaped to press the lever for a reward. To avoid overtraining, sessions were terminated after 10 lever presses. Lever training was complete when rats met the 10-press criterion on each side of the maze. Habituation and lever training was completed before rats underwent surgery.

Following recovery from surgery, subjects received two vertical exploration sessions, one in each vertical column (Figure 2). Previous research confirms that rats navigate space in the vertical plane (Grieves et al., 2020), and prior data from our lab suggests that rats best attend to demonstrators in the VM if first exploring vertical columns (Wise et al., 2024). For vertical exploration, empty holding cages were stacked in each vertical column. Subjects entered the stacked cages through a hatch door in the cage furthest from the test platform where they could then travel up and down the vertical column. Access holes on the floor and lid of each cage as well as a ramp system were used to facilitate vertical exploration. A final access hole in the testing platform allowed rats to explore the entirety of the maze within a session. Precision pellets were scattered throughout the holding cages, testing platform, and pellet receptacles to encourage exploration. Criteria for completing vertical exploration was that rats must travel from the bottom holding cage to the test platform and back for both the West and East sides of the maze. All subjects met criteria within 2-3 sessions and no adverse reactions to the apparatus were observed.

#### Spatial Distance Training

One to three days prior to spatial distance training, rats received one lever pressing refresher session per test chamber. The first phase of training required that subjects learn to discriminate demonstrators based on spatial distance (Figure 2).

Spatial distance included demonstrators placed in the farthest vertical position from the testing platform, referred to as “far-spatial”, or in the closest vertical position, referred to as “near-spatial”. Half of the subjects in each surgery group were in the far-spatial condition (rewarded for choosing the farthest demonstrator) and half were in the near- spatial condition (rewarded for choosing the nearest demonstrator) for the duration of spatial training. For each subject, two equally unfamiliar demonstrators were used in each trial for the duration of training. Demonstrator position was pseudo-randomly organized so that each demonstrator rat was assigned as both far- and near-spatial and appeared in both the West and East columns for an equal number of trials per session. Counter-balancing distance and side ensured that spatial distance was the only relevant dimension.

Spatial training consisted of 20 trials per session for 12 daily sessions. Before each trial subjects were placed in the closed center chamber and demonstrators were positioned at the near and far vertical distances. At the start of each trial, guillotine doors were lifted, lights were turned on, and levers were extended in the West and East chambers. Each chamber included a wire grid floor that allowed subjects to perceive demonstrators positioned below. Choice was indicated by a lever press in either the West or East chamber. Correct choices were signaled by a brief tone, and three pellets were dispensed before all lights were turned off and levers were retracted. Each session began with five discovery trials. If a subject’s first choice was incorrect, the pressed lever was retracted, and the correct lever remained extended until pressed, resulting in the tone and delivery of 3 pellets. Discovery trials ensured that spatial distance discriminations were reinforced before subjects continued testing. After 5 discovery trials, an incorrect choice terminated the trial. At the end of each trial, subjects were placed back in the center chamber with doors closed and demonstrators were positioned for the next trial. Demonstrators were assigned individual holding cages to limit potential cross-odor contamination. Test chambers and vertical columns were cleaned and aired out with a handheld fan between subjects to reduce odor cues. Once spatial training was complete, subjects began domain transfer testing the following day.

#### Domain Transfer Testing

Domain transfer testing began immediately after training. Each of the four test sessions consisted of 16 spatial distance trials and 4 social distance trials, otherwise known as domain transfer probes. Domain transfer probes were pseudo-randomly presented so that they were at least three trials apart and differently ordered for the first and second inactivation sessions. General procedures remained consistent across spatial training and domain transfer testing, including experimental set-up, counterbalancing, and trials per session. In the four domain transfer probes, one of the two demonstrators was replaced with the subject’s cagemate, and both demonstrator and cagemate were placed at the same vertical distance level (Figure 2C). The demonstrator used was counterbalanced across probe trials. The demonstrator and cagemate were each placed in the middle position of one of the two vertical columns, therefore removing any difference in spatial distance. Instead, non-subject rats now differed in social distance, i.e. relative familiarity. Placement of non-subject rats in the West and East columns was counterbalanced across probe trials. In each probe trial, subjects were allowed to select either the demonstrator or the cagemate via lever press, and no reward was given for either choice. Correct spatial-to-social domain transfer was defined as when a social distance of the selected rat aligned with the previously rewarded spatial distance, i.e. a rat in the near-spatial training condition would select the cagemate and a rat in the far-spatial training condition would select the less familiar demonstrator. To examine whether the PPC was required for domain transfer, chemogenetic inactivation of the area preceded domain transfer testing for all rats in half of the sessions. Clozapine n-oxide (CNO, 5 mg/kg) was injected for probe sessions 1 and 3 of all rats. Saline was injected in probe sessions 2 and 4. Approximately 40-50 minutes before testing, subjects received an intraperitoneal injection of either CNO or saline and were returned to their home cage. CNO was not counterbalanced within sessions because of the concern that animals might learn that certain trials would not be reinforced. In that case the probe trials in the first session would be most informative.

### Analysis

Analyses were conducted on all subjects for spatial training and domain transfer testing. Spatial training data on sham animals only was also reported in an earlier methods paper (Wise et al., 2024). Response accuracy and latency were recorded for all test sessions. Trials were considered correct when subjects pressed the lever positioned above the vertical column that held the correct spatially or socially distanced demonstrator. For discovery trials, choice was considered correct when subjects pressed the appropriate lever on the first try. Latency was defined as the duration between trial start and lever press. For spatial training, accuracy and latency were analyzed using a repeated measures analysis of variance (RMANOVA) with surgery group as the between-subjects variable and session as the within-subject variable.

Control analyses were conducted to assess whether there were differences between the near- and far-spatial training conditions. For domain transfer testing, accuracy and latency response were again analyzed using a RMANOVA. Tests for spatial (non-probe) and domain transfer probe trials were run separately. Again, control analyses were conducted to assess whether there were differences between the near- and far-spatial training conditions. Finally, a binomial distribution was calculated to determine the probability that the proportion of correct spatial-to-social transfer choices for each session was above chance.

### Histology

Following the completion of behavioral testing, subjects were deeply anesthetized with isoflurane and given an intraperitoneal overdose of sodium pentobarbital (Beauthansia-D; 100 mg/kg). Rats were perfused transcardinally first with phosphate-buffered saline and then with 10% (w/v) formalin in a 0.1m phosphate buffer. Brains were post-fixed for at least 48 hours in the 10% formalin solution at 4° C before being cryoprotected in 30% (w/v) sucrose solution for at least 48 hours. Next, brains were sectioned coronally into three series of 40 μm sections using a microtome. Series were stained with thionine for Nissl material, DAPI for fluorescence, or kept as a spare.

Nissl images were captured to compare against fluorescent images in estimating bregma. Fluorescent images were captured in the blue channel for DAPI-stained neurons and in the red channel for neurons transfected by the DREADDs virus. Fluorescent images were used to evaluate expression in the targeted PPC areas as well as extra-target spread. Cases were considered acceptable when fluorescence was distributed bilaterally in the dorsal PPC and when extra-target damage was not bilateral and/or was not widely distributed in the any extra-target region.

## RESULTS

### Histology

Viral transduction was observed bilaterally for all PPC surgery rats (Figure 3). In one case, there was unilateral, partial involvement of primary somatosensory cortex. Thus, no rats were excluded based on histology.

**Figure 3.**
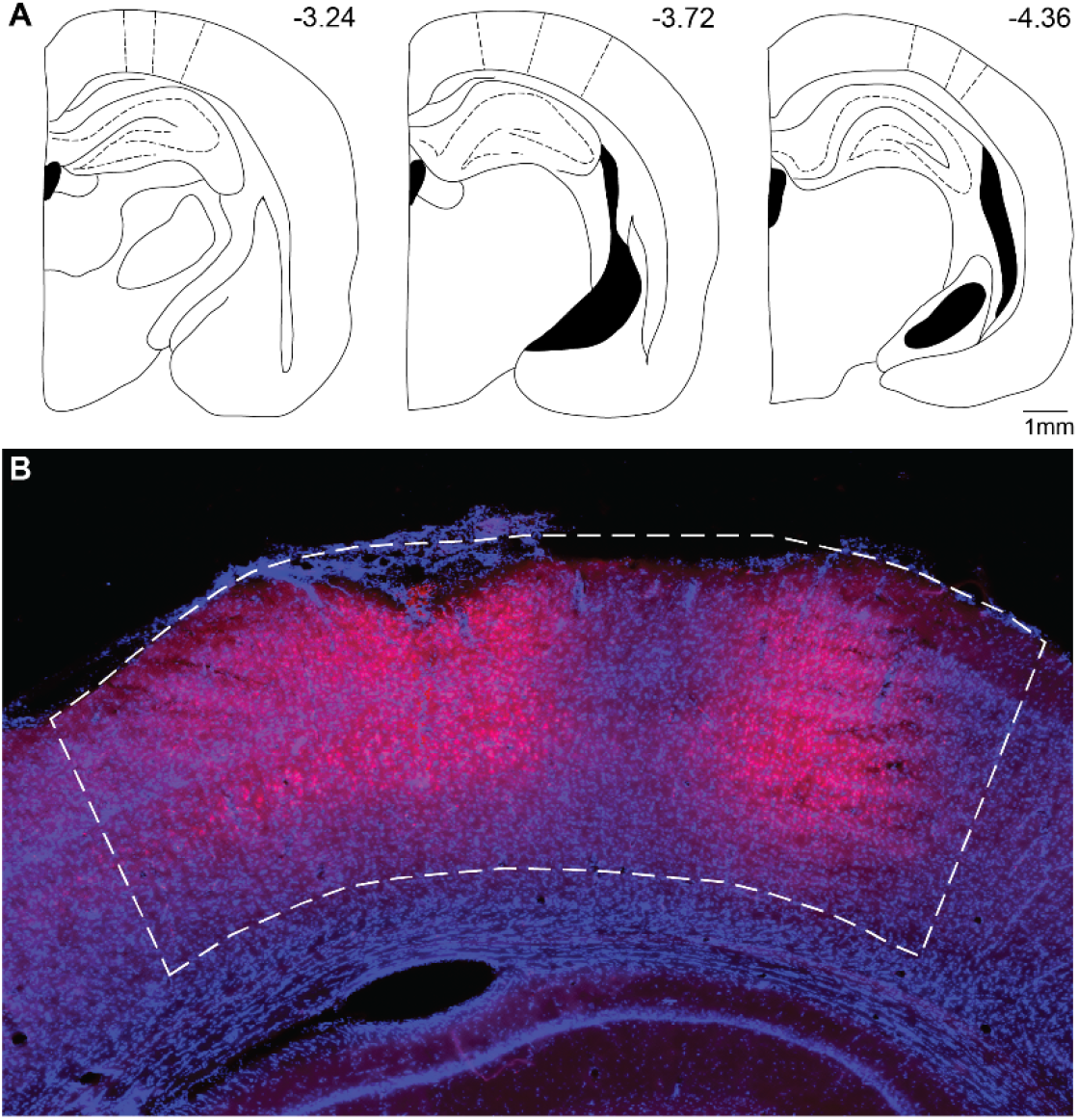
Viral injection site. Representative schematics and viral transduction labeling of the dorsal PPC. A) Three schematics of the PPC across bregma levels. PPC borders illustrated by dashed lines. B) Representative viral transduction within PPC borders at bregma -3.60.

### Behavior

#### Spatial Distance Training

Prior to training all rats met lever pressing criteria in each of the outer chambers within 2-3 sessions (4-6 sessions total). In spatial distance training all rats demonstrated improved performance across trials (Figure 4, middle) confirmed by a main effect of session (F_1,11_ = 5.895, p < 0.001), but no effect of group (p = 0.571) or ‘session x group’ interaction (p = 0.821). Likewise, latency to make a choice also decreased with training (Figure 4, right) and was confirmed by a main effect of session (F_1,11_ = 20.125, p < 0.001), but no main effect of group (p = 0.270) or ‘session x group’ interaction (p = 0.152). No group effects were expected given that the PPC was not inactivated.

**Figure 4.**
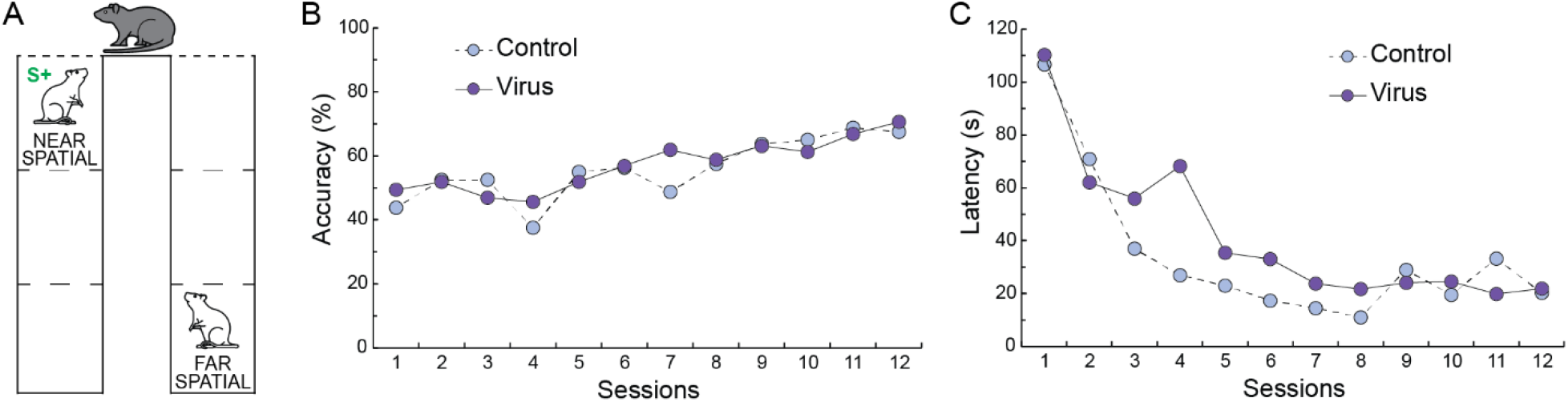
Training Results. A) Training required that subjects learn to select either the near- (pictured) or far-spatial distance of a conspecific. Correct choice was based exclusively on the spatial dimension. B) Accuracy increased over time for both PPC virus and control groups. Both groups were performing at approximately 70% correct at the completion of training. C) Response latency decreased over time for both groups. S+ indicates rewarded choice.

Additionally, there were no differences between training condition (near-spatial vs far- spatial) for either accuracy (p = 0.300) or latency (p = 0.243). Together, these results suggest that rats effectively learned to discriminate between the spatial distance of two conspecifics in a goal-direct paradigm whether the correct choice was the near conspecific or the far conspecific.

#### The Role of the PPC in Domain Transfer

Each domain transfer testing session consisted of 16 spatial distance trials and four domain transfer (social distance) probes presented in a pseudorandom order.

Spatial and social distance were analyzed separately. For these analyses, group was the between-subject variable, and drug condition (CNO vs SAL) and drug condition session (first vs second) were the within subject variables. We conducted the same analyses for training condition. A group by drug condition interaction would suggest that the PPC is involved in spatial-to-social information transfer.

For spatial distance accuracy, groups performed similarly on the first CNO and SAL days, but on the second CNO and SAL days, the control group showed increased accuracy and the virus group showed decreased (CNO) or the same (SAL) accuracy (Figure 5). This was confirmed by a significant group x session interaction (F_3,10_ = 7.832, p = 0.019) to each other and across the four sessions. There were no main effects of group, drug condition, or session and no other significant interactions (p > 0.129). For latency on spatial distance trials, the control group demonstrated higher latencies than the virus on the first and second SAL days and on the second CNO day (Figure 5). This was confirmed by two significant interactions: ‘drug condition x session’ (F_1,10_ = 7.343, p = 0.022) and ‘group x drug condition x session’ (F_3,10_ = 6.069, p = 0.033). Otherwise, the ‘group x session’ interaction was not significant (p = 0.886), but all other main effects and interactions tests there were a number of marginally significant effects (p ranged from 0.051 to 0.094). Although there were hints of a ‘group x drug condition’ interaction for latencies, the effects were not in the direction that would provide evidence for involvement of the PPC.

**Figure 5.**
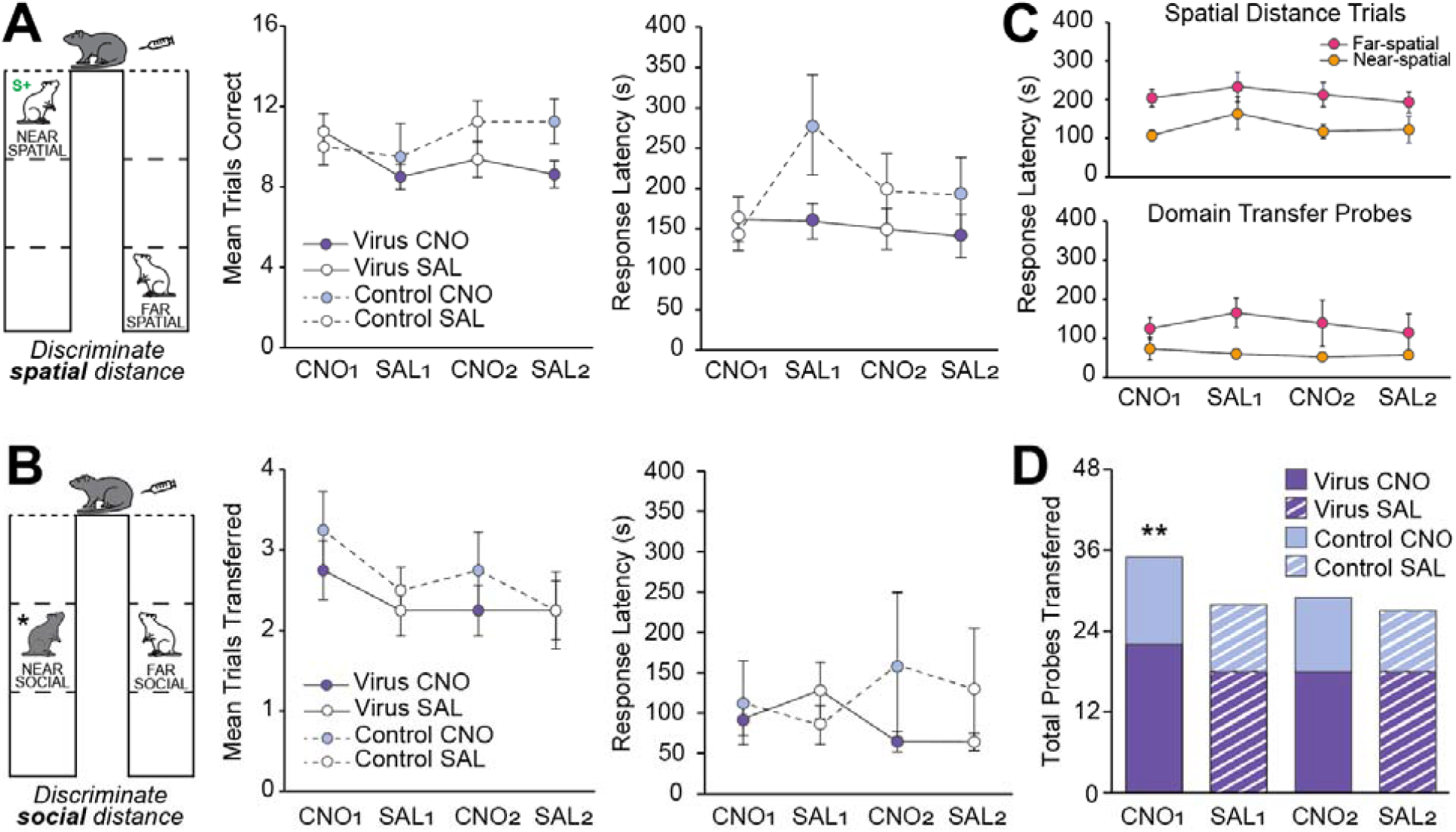
Domain transfer results. Injections of either CNO or saline (SAL) were administered prior to testing on each session. One session was run per day for four consecutive days. A) Accuracy and latency results for spatial distance trials of domain transfer testing. Accuracy was similar for PPC virus and controls groups on the first CNO and SAL days but was increased for controls and either decreased or the same for the virus group on the second CNO and SAL days. Response latency was higher for the control group as compared to the virus group on the first and second SAL day and first CNO day. B) Accuracy and latency results for the domain transfer probes. No differences in accuracy or latency were found between PPPC virus and control groups. C) Latency results compared across training condition (near-spatial vs far-spatial). Rats trained in the far-spatial condition showed higher latencies in both spatial distance trials (top) and domain transfer probes (bottom) across CNO and SAL sessions. D) Total domain transfer probes correct for each domain transfer testing session. Cumulative probability for total probes correct was significantly above chance for the first testing session. PPC inactivation did not impact number of correct domain transfer probes. ** indicates p < 0.01.

Training condition (near-spatial vs far-spatial) had little effect on spatial distance accuracy. Although the ‘drug condition x session’ interaction was marginally significant (p = 0.075), all other tests were not significant (p > 0.215). Training condition did, however, impact latency such that animals trained in the far-spatial condition showed higher latencies (Figure 5C). This was confirmed by a significant effect of training condition (F_1,10_ = 5.768, p=0.037) and session (F_1,10_ = 5.287, p = 0.044).

For social distance accuracy, the control and virus groups performed similarly across drug conditions and sessions (Figure 5). There were no main effects of group, drug condition, or session and no significant interactions (p > 0.240). Regarding latencies, the two groups were similar on the first CNO day, but the control group tended to show higher latencies than the virus group on the first SAL day (Figure 5). There was a marginally significant ‘group x session’ interaction (F_1,10_ = 4.029, p = 0.072). All other main effects and interactions were not significant (p > 0.203). There was no evidence for involvement of the PPC.

Training condition had little effect on social distance accuracy. Although the ‘drug condition x distance’ interaction was marginally significant (p = 0.098), all other tests were not significant (p > 0.234). There was a tendency for training condition to impact latency such that animals trained in the far-spatial condition showed higher latencies, especially on the first SAL session (Figure 5C). Accordingly, there was a marginally significant effect of training condition (p = 0.057). All other tests for latency were not significant (p > 0.154).

#### Evidence for Spatial to Social Domain Transfer

An important goal of the study was to assess whether there was evidence that rats transfer spatial distance information to the social distance domain. Given the lack of evidence for involvement of the PPC in domain transfer testing, the total number of correct transfer probes across all rats was examined. Probe sessions were analyzed separately because of the possibility that rats might learn that there were unrewarded trials. In session 1, rats selected the appropriate social distance in 35 out of 48 total domain transfer probes. Binomial analysis indicates that the cumulative probability of this happening by chance is P(X≥35) < 0.001. In subsequent sessions the cumulative probabilities were 28 of 48, 29 of 48, and 27 of 48 choices. Binomial analysis indicates that the cumulative probability of choosing 27 out of 48 by chance is P(X≥27) > 0.235.

## DISCUSSION

In this study we investigated three main questions: 1) can rats learn to discriminate differences in spatial distance from two conspecifics, 2) do rats transfer learned spatial distance information to the social distance domain, and 3) does the PPC play a role in spatial-to-social domain transfer. We provide evidence that rats can learn to discriminate spatial distances of conspecifics for a reward and then transfer spatial information to the social distance domain. There was no evidence that PPC inactivation impacts spatial-to-social domain transfer.

Rats readily learned to discriminate conspecifics at either the near or far spatial distance using only the spatial dimension. These findings are consistent with prior research showing that rats discriminate spatial information in unstructured paradigms (Goodrich-Hunsaker et al., 2005; Save et al., 1992; Warburton & Brown, 2015) and can maintain goal-driven discriminations over time (Fox, Barense, & Baxter, 2003; Nitz, 2006). Possibly due to depth perception limitations and image size confounds, goal-oriented spatial distance discrimination has not been reported outside of this study. We addressed these limitations by presenting information-rich stimuli in a novel apparatus designed for multisensory perception and discrimination. One key for successful discrimination in the VM is allowing rats to explore the entire maze prior to testing in order to map vertical space (Grieves et al., 2020; Wise et al., 2024). We show that just as rats are highly adept at detecting spatial differences through spontaneous exploration, they are also skilled in differentiating spatial distance in a goal-oriented manner. These findings reveal a novel capacity for rats’ spatial cognition beyond spontaneous exploration and other goal-driven spatial behavior.

Upon learning to discriminate spatial distance between conspecifics, rats underwent domain transfer testing. For most trials, rats were presented with two equally familiar demonstrators at separate spatial distances. Both groups correctly discriminated spatial distance, and accuracy did not differ between subjects trained to select the near and far spatial distance. Rats in the far training group showed longer latencies to respond than the near training group for both spatial distance trials and domain transfer probes. CNO dosages were identical across training groups, making it unlikely that off-target behavioral effects associated with the drug (MacLaren et al., 2016) explain latency differences. Further, longer latencies for the far training group were observed across CNO and SAL sessions. Another possibility is that altered task expectations following the introduction of domain transfer probes slowed performance. Rats will increase exploration following object-in-context transformations (Dix & Aggleton, 1999; Sep, Vellinga, Sarabdjitsingh, & Joëls, 2021), so similar behaviors may be expected when subjects experience their cagemate in the VM. However, this does not explain longer latencies specific to the far training group. Alternatively, it may be the case that accurately identifying a demonstrator at the far spatial distance is either perceptually or cognitively more difficult. Yet, latency differences were only observed in domain transfer testing and not in earlier training. A more comprehensive explanation which includes both theories is that latency results were due to an interaction between changing task expectations and an increased perceptual or cognitive load on the far training group. Given that accuracy did not differ between training groups, this theory suggests that rats overcome potential difficulties in perceiving conspecifics at a far- spatial distance even when expected task demands become inconsistent.

Domain transfer testing included a subset of probe trials for testing rats’ ability to transfer learned spatial knowledge to a social context. Domain transfer probes removed all differences in spatial distance while creating a difference in social distance. When initially challenged to discriminate social distance, rats were correct for a significant majority of probe trials. To our knowledge, this is the first evidence that rats transfer information across spatial and social cognitive domains. This provides support for high levels of spatial abstraction. Probe accuracy declined in subsequent sessions but was still numerically above chance. Subjects may have learned that social probe trials were never reinforced. Rats can adaptively respond to reward omission by pivoting to explore reward opportunities in other locations (Papini, Guarino, Hagen, & Torres, 2022). Near vs far training condition had no impact on evidence for domain transfer, suggesting that spatial to social information transfer does not rely on specific distances. This aligns with cognitive mapping theories that ego-centric social distance of those both closer and farther away are used to construct abstract social space. For our study, rats’ ability to accurately match social distance to spatial distance, regardless of initially learned distance, hints at a general framework for processing spatial and social distance while providing robust evidence for domain transfer.

Our results provide novel insights into rats’ flexible decision making and attention in spatial and social contexts. In goal-oriented tasks, rats can direct attention towards specific stimulus dimensions while disregarding others (Birrell & Brown, 2000; Bucci, 2009; Tait, Chase, & Brown, 2014). Notably, rats adaptively discriminate stimuli in a modified version of the Wisconsin card sorting task, known as the attentional set shifting task (AST; (Birrell & Brown, 2000; Tait et al., 2014). AST incorporates multidimensional stimuli (digging medium and odor) with only one dimension relevant to obtaining a reward. Our study also tasked rats at discriminating and shifting attention between relevant stimuli dimensions, albeit spatial and social dimensions. Domain transfer testing may be comparable to interdimensional (ID) shifts in AST where rats are quick to shift attention to dimensions that were once irrelevant but now inform correct choice (Tait et al., 2018; Tait et al., 2014). Consistent with AST results, our research adds that attentional shifting behavior applies to spatial and social dimensions of information-rich stimuli. We also found that domain transfer performance did not depend on whether correct social distance was defined as the cagemate or less familiar demonstrator. This suggests that not only can rats use social identity as a relevant feature for goal-driven decision making, but also that inherent social preferences do not universally dictate rats’ behavior in social contexts. Under both training and test conditions subjects may have needed to override innate social preferences to complete the task. This is supported by findings from our lab that rats can disregard competing social novelty preferences to select previously reinforced conspecifics (Mazumder, 2023). Other research shows that rats, via lever press, will select to interact with a less familiar rat over their cagemate (Hackenberg et al., 2021), suggesting that near-trained subjects in our study likely overrode social preferences to complete domain transfer probes. Substantial literature confirms social novelty preference in rats (Acikgoz et al., 2022; Beery & Shambaugh, 2021; Hackenberg et al., 2021; Templer et al., 2018), however to our knowledge our study is the first to report that rats can use social identity to inform decisions that may go against spontaneous social preference. Social discrimination between two conspecifics for a food reward has been reported elsewhere only once before (Gheusi et al., 1997). Rats’ ability to adaptively attend to spatial and social information in this study shows promise that these animals can be used as models for complex decision making in both domains.

We found no evidence that the PPC was required for spatial distance discrimination or domain transfer. Impairments in spatial discrimination were not expected given that the PPC contributes to processing spatial changes (Goodrich-Hunsaker, Howard, Hunsaker, & Kesner, 2008; Goodrich-Hunsaker et al., 2005; Save et al., 1992) but not acquisition (Scott et al., 2021). To ensure adequate task knowledge before domain transfer the PPC was not inactivated during training. Therefore, knowledge to complete spatial trials during domain transfer testing (with PPC inactivation) had already been acquired. Spared spatial performance is consistent with prior findings that PPC lesioned rats still detect changes in spatial distance between landmarks (Goodrich-Hunsaker et al., 2005) and discriminate novel spatial locations (Wise et al., in press); though goal-driven spatial discrimination has not been reported outside of our study. Unexpectedly, PPC inactivation did not lead to performance deficits during domain transfer. CNO injections may have been insufficient in inactivating the PPC. However, multisensory decision-making impairments in rats with PPC inactivation have been reported with CNO dosages lower than our own (Raposo, Kaufman, & Churchland, 2014). A broader explanation is that domain transfer testing did not tap into complex decision making processes attributed to the PPC. The PPC is engaged under complex spatial demands (Scott et al., 2021), but whether spatial-social discriminations fall under this umbrella are unknown. In non-spatial discriminations such as ID shifts during AST, PPC deficits do not impair performance (Bucci, 2009; Fox et al., 2003).

Perhaps the same is true when rats are tasked at shifting from spatial to social dimensions. Despite evidence that the PPC is implicated in separately estimating spatial and social distance in humans (Yamakawa et al., 2009) and integrating multisensory information in multiple species (Andersen, 1997; Freedman & Ibos, 2018; Mohan, de Haan, Mansvelder, & de Kock, 2018), it may not be required for spatial-to-social transformations. Alternatively, virus animals in our study may have overcome PPC deficits by recruiting different cortical regions to transfer information. Bypassing PPC dysfunction by relying on other brain regions aligns with cognitive mapping research demonstrating that abstract processing employs a broad network of cortical areas, including the hippocampal formation (Montagrin et al., 2018; Schafer & Schiller, 2018; Tavares et al., 2015) and orbital frontal cortex (Qiu et al., 2024). Our results suggest that the PPC is not required for spatial-social domain transfer, but we cannot conclude whether the region partially contributes to their shared mechanisms. Indeed, the PPC is implicated in a much broader cortical network for complex decision-making and multisensory integration. More research is needed to determine PPC engagement during domain transfer in rats and other species. Future studies investigating the role of the PPC in spatial-social cognition may benefit from employing *in vivo* recording techniques capable of capturing PPC activity and/or circuit analyses for PPC-HP connections.

## CONCLUSION

Our study concludes that under complex task demands rats can learn to discriminate spatial distances of social stimuli and later transfer learn information onto a social context. This is the first reported evidence of spatial-social domain transfer in rats. In line with cognitive mapping and spatial abstraction theories, our results show that spatial and social distance processing in rats may rely on shared spatial-social mechanisms. While prior evidence shows that the PPC is implicated in spatial and social processing, results from our study suggest that this region is not be required for domain transfer. Using a novel paradigm informed by species-specific perception, our results highlight and expand on the complex decision making capabilities of rats that reach beyond known spatial and social behavior.

## Acknowledgements

We would like to thank John Murphy for his assistance in engineering, constructing, and refining the Vertical Maze. We would also like to thank Isabella Pilkington and Jose Pena for their assistance in data collection as well as Xiangyuan Peng, Ph.D. and Devon Poeta, M.S. for their advice. This work was funded by NSF IOS 1656488 and NIMH R01MH108729 to RDB and NINDS F99 NS129180-01A1 to TBW. TBW is now located at the Department of Psychology, Yale University, New Haven, CT 06510, USA.

